# Mathematical expressions describing enzyme velocity and inhibition at high enzyme concentration

**DOI:** 10.1101/2021.12.05.471285

**Authors:** Agustin Hernandez

## Abstract

Enzyme behaviour is typically characterised in the laboratory using very diluted solutions of enzyme. However, *in vivo* processes usually occur at [S_T_] ≈ [E_T_] ≈ K_m_. Furthermore, the study of enzyme action usually involves analysis and characterisation of inhibitors and their mechanisms. However, to date, there have been no reports proposing mathematical expressions that can be used to describe enzyme activity at high enzyme concentration apart from the simplest single substrate, irreversible case. Using a continued fraction approach, equations can be easily derived to apply to the most common cases in monosubstrate reactions, such as irreversible or reversible reactions and small molecule (inhibitor or activator) kinetic interactions. These expressions are simple and can be understood as an extension of the classical Michaelis-Menten equations. A first analysis of these expressions permits to deduce some differences at high *vs* low enzyme concentration, such as the greater effectiveness of allosteric inhibitors compared to catalytic ones. Also, they can be used to understand catalyst saturation in a reaction. Although they can be linearised following classical approaches, these equations also show some differences that need to be taken into account. The most important one may be the different meaning of line intersection points in Dixon plots. All in all, these expressions may be useful tools for the translation *in vivo* of *in vitro* experimental data or for modelling *in vivo* and biotechnological processes.

## INTRODUCTION

Enzymes in the laboratory are usually assayed in conditions that are considered close to the *in vivo* conditions but, actually, they may differ significantly in some aspects from those governing their action in the cell. This includes variations in pH, crowding and ionic strength, among others. Particularly, it is very frequent to observe that, *in vivo*, enzymes are catalysing reactions in conditions where the concentration of substrate is close to that of the enzyme. On the contrary, in the laboratory, a typical enzyme assay is done in conditions where the concentration of the enzyme is as low as possible. This difference is a major one to understand the real activity of enzymes *in vivo*, because although it is usually assumed there exists a linear relationship between enzyme concentration and activity [1] that is not necessarily the case. The usual approaches for prediction of the initial velocity of an enzyme reaction assume either a fast substrate-enzyme binding equilibrium (FE), as originally proposed by Michaelis and Menten [2] or, more often, a quasi-steady state for the variation in the concentration of the enzyme-substrate complex (QSS), as the modification introduced by Briggs and Haldane [3]. In addition, both approaches assume that the concentration of the enzyme is negligible compared to that of the substrate. This is an important assumption when deriving the rate expressions and the one that dictates how enzyme assays are done in the laboratory. Several mathematical approaches have been proposed to derive expressions that could be valid in conditions where the concentration of the enzyme and substrate are close [4–7]. Moreover, a so-called inverse Michaelis-Menten expression has been proposed for interfacial enzyme kinetics [8]. However, to the best of this researcher’s knowledge, none yet have dealt with situations different from the simplest irreversible single substrate reaction.

In addition to an interest in describing the biochemistry of living organisms unperturbed, in pharmacology and biochemistry it is especially important the capacity to describe the effect of small molecules on enzyme activity, such as inhibitors and activators. Again, small molecules are assayed and characterised using low enzyme concentrations, but since target enzymes in the cell are in much closer ratios with their substrates, the effect of those small molecules *in vivo* may depart from the behaviour found *in vitro*.

From the point of view of the experimental biochemist, the mathematical expressions describing enzyme action *in vivo*, or in a biotechnological setting, should ideally use parameters that can be easily estimated in the laboratory. In addition, the possibility of creating plots that can show differences between assay or living conditions can help in the interpretation of results. From the point of view of teaching biochemistry, the use of uncomplicated algebra and calculus is a bonus when explaining the derivation of those expressions.

In this study, I present a series of mathematical expressions useful to describe the velocity of monosubstrate enzyme reactions in different conditions when the concentration of the enzyme is near that of the substrate. Those can be used with estimates of parameters obtained in the laboratory at low enzyme-to-substrate concentration ratios to obtain biologically relevant conclusions.

## METHODS

Let be [S], [P] and [E] the respective free concentrations of substrate, product and enzyme in the reaction depicted in Scheme 1. The microscopic kinetic constants are numbered using odd figures for those representing forward reactions with respect to product formation, while reverse reactions receive corresponding even numbering. Initial conditions for the reaction (t=0) are assumed and, therefore, [P] ≈ 0 is also assumed, leading to irreversibility of the reaction. Although incorrect, for simplicity’s sake only, that last assumption is depicted by the corresponding microscopic kinetic constant as showing zero value in the scheme. The presence of high concentrations of enzyme has been approached by considering that the following mass conservation expression needs to be taken into account: [S_T_] = [S] + [ES], where [S_T_] stands for total amount of substrate in the reaction. In the case of a reversible reaction, [P_T_] = [P] + [EP] was also contemplated. These conservation laws are considered in addition to the usual expression for the conservation of enzyme, [E_T_] = [E] + [ES], where [E_T_] stands for the total amount of enzyme in the system. Assuming QSS conditions for the concentration of the enzyme-substrate complex ([ES]), the following expression can be derived (a detailed process of derivation is shown in Supplemental Material 1A):

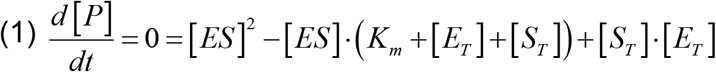

**Scheme 1.**
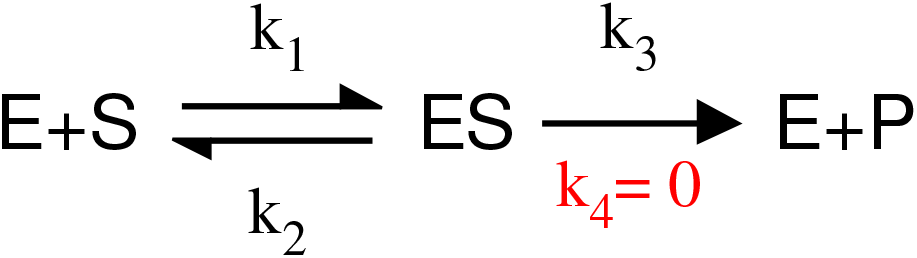

Solving this quadratic expression by regular means leads to

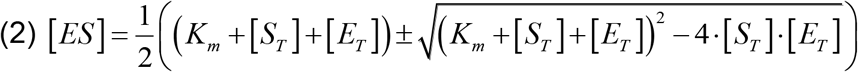

However, a simpler approach to approximate (1) can be taken by using continued fractions (CF). This method uses simple algebra and has been used extensively in other areas [9, 10]. In general, a quadratic expression where the coefficient for the quadratic term is one, i.e. an expression of the type:

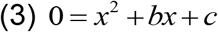

can be approximated for both negative and positive sign solutions as:

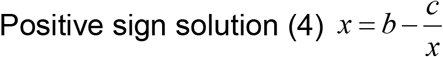

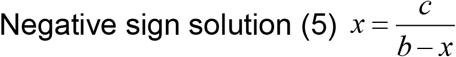

Recursive substitution of the variable *x* in the left side of the equality with the whole expression on that same side permits increasing degrees of approximation (for an example, see Supplemental Material 1B and Figure 1A). A first degree approximant can be obtained by neglecting the variable *x* in the left hand side of the equality. In this study, first degree approximants were used only. On the other hand, the negative sign solution of (2) is the only one valid in chemistry. This can be exemplified by the fact that, in the case of the positive sign, when [S_T_] = 0, the value of [ES] is non-zero:

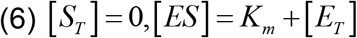

**Figure 1.**
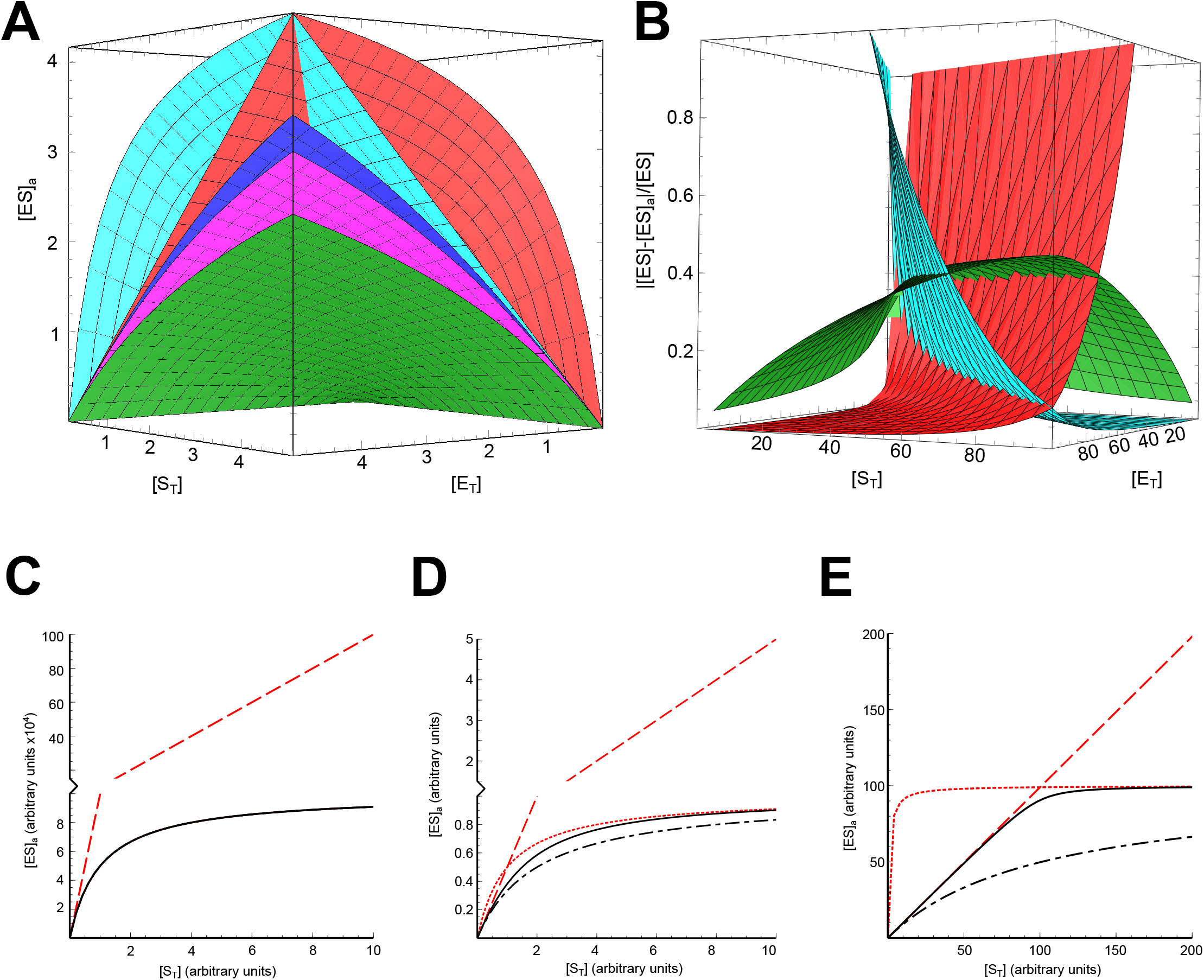
Estimation of enzyme-substrate complex concentration using different mathematical expressions. **A** Approximated [ES] ([ES]_a_) using the exact solution (expression (2) in main text; blue surface), extended Michaelis-Menten (eMM; expression (7) in main text; green surface), a second degree approximant (eMM, expression (19) from Supplemental Material 1; magenta surface), standard Michaelis-Menten (expression (7b) in main text, cyan surface) and inversed Michaelis-Menten (expression (7c) in main text; red surface). All surfaces calculated using K_m_ = 1. **B** Estimated error relative to the real value estimated with the exact solution and the extended Michaelis-Menten (eMM; expression (7); green surface), standard Michaelis-Menten (expression (7b); cyan surface) and inversed Michaelis-Menten (expression (7c); red surface). All surfaces calculated using K_m_ = 1. **C** Departure of [ES]_a_ from the exact solution at a constant low enzyme concentration ([E_T_] =0.001 *x* K_m_). Solid black line: exact solution (expression (2) in main text); Dot-dashed black line: extended Michaelis-Menten (eMM; expression (7)); Dotted red line: standard Michaelis-Menten (expression (7b)); Dashed red line: inversed Michaelis-Menten (expression (7c)). **D** Departure of [ES]_a_ from the exact solution at a constant enzyme concentration similar to K_m_ ([E_T_] = K_m_). Line identities as in C. **E** Departure of [ES]_a_ from the exact solution at a constant high enzyme concentration ([E_T_] = 100 *x* K_m_). Line identities as in C. All lines in C, D, and E calculated using K_m_ = 1. Please note that eMM and sMM lines in panel C are obscured by the overlapping line corresponding to the exact solution.

The CF approach for the negative sign solution has been used with (1) and all other reaction schemes treated in this study. Detailed derivation of the expressions can be found in Supplemental Material 1 and 2.

## RESULTS

### Derivation of expressions for monosubstrate irreversible reactions

The CF approximation to solving [ES] in (1) under QSS assumption leads to the expression (Supplemental Material 1, B):

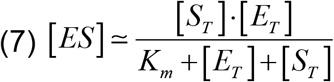

Expression (7) can be reduced to either that obtained using standard Michaelis-Menten assumptions (sMM) [3]

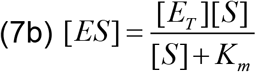

or the inverse Michaelis-Menten (iMM) [8]

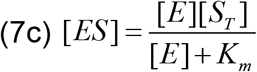

in cases where [E_T_] ≪ [S_T_] or [E_T_] ≫ [S_T_], respectively. Therefore, (7) can be understood as an extended Michaelis-Menten (eMM) bridging two extremes. Indeed, a comparison between the values estimated for [ES] using one as a token value for K_m_ show that expression (7) produces approximate values of [ES] ([ES]_a_) that show a similar behaviour to the real value throughout the range of [E_T_] and [S_T_] values (Figure 1A), although they underestimate the true value of [ES]. Greater accuracy can be obtained by higher degree approximants or a wise use of the sMM and iMM (Figure 1A and 1B). Nevertheless, a first degree approximant already provides a much closer approximation to the value of [ES] than sMM or iMM approaches when the whole range of [S_T_] and [E_T_] is taken into account (Figure 1A and 1B). The relative value of the error for (7) was found never to exceed 0.4 (Figure 1B), while that for sMM and iMM increased exponentially as conditions departed from the assumed ones for their derivation.

When [E_T_] was assumed constant at a very low concentration (e.g. [E_T_] = 0.001 ·K_m_), it could be observed that [ES]_a_ from both sMM and eMM produced curves that overlapped the values obtained using the exact solution (expression (2)) over the range 0 ≥ [S_T_] ≤ 10·K_m_, while iMM generated gross overestimations (Figure 1C). As [E_T_] is increased, the curves generated by sMM and eMM separated from the true value of [ES]. For example, if 1 = [E_T_] = K_m_, sMM produced moderate overestimations while eMM underestimated the value of [ES] (Figure 1D). These deviations tended to asymptotically converge with the true value of [ES] at high [S_T_]. This behaviour was exacerbated if greater [E_T_] were contemplated (Figure 1E). A similar phenomenon can be observed if [S_T_] is kept constant and curves are created over ranges of [E_T_] similar to those used in Figures 1C to 1E for [S_T_] (data not shown). However, in that case, sMM is the expression providing gross overestimations, while iMM produces moderate overestimations, and eMM keeps on producing moderate underestimations of the true value of [ES].

QSS assumption may not be appropriate in all cases. Nevertheless, the CF approach can also be used using FE assumptions and, in the present case, leads to the similar expression (supplemental material 1C):

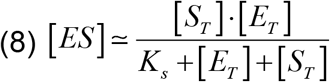

Expressions (7) and (8) can be used to provide rate laws for a monosubstrate irreversible reaction under the conditions stated (Table 1). In particular, under QSS assumption:

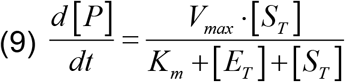

As it is easily observed in (9), the catalytic efficiency, i.e. the value of the kinetic constant for the reaction when it shows first order kinetics, is not V_max_/K_m_ but V_max_/(K_m_+[E_T_]). This also leads to the observation that K_m_ does not coincide with either the concentration of substrate providing 1/2V_max_ or the abscissa value of the intersection point between the lines describing the velocity of the reaction when first order and when zero order. In both cases, it is easily demonstrated that the value for both is K_m_+[E_T_ (Figure 2A).

**Figure 2.**
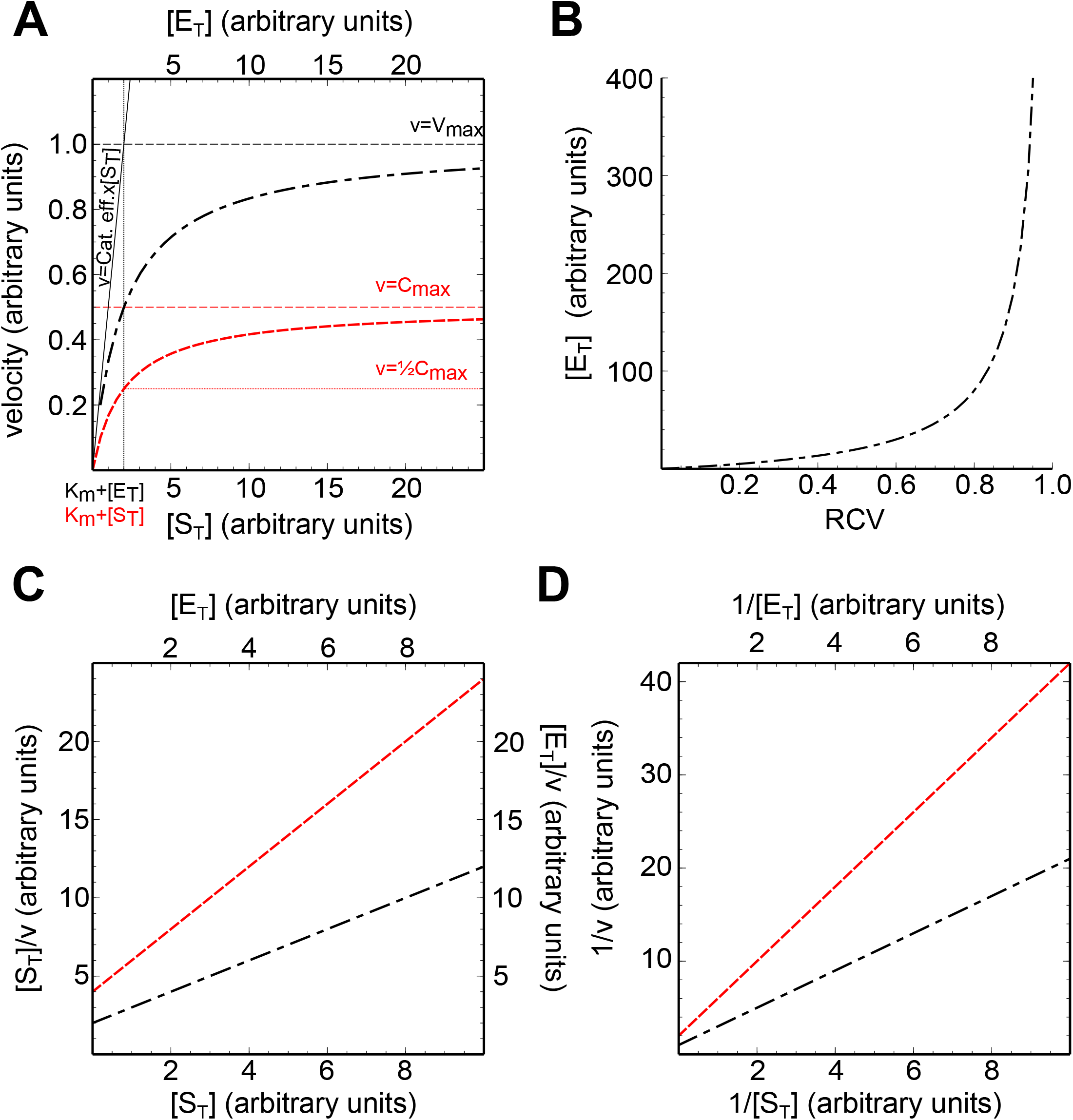
Hyperbolic and linear plots of eMM expressions for monosubstrate irreversible reactions. **A**. Direct plots of the estimated velocities (equation (9)) at constant [E_T_] (Dot-dashed black line) or constant [S_T_] (Dashed red line). Both lines calculated using K_m_ = 1. Value for the concentration of the constant reactant was set to one (1). To help visualisation, *k_3_* = 1 was used for constant [E_T_] while *k_3_* = 0.5 was used for constant [S_T_]. Other parameter values: [E_T_] = [S_T_] = K_m_=1. Cat. eff.: catalytic efficiency. **B** Concentration of [E_T_] as a function of Relative Catalytic Velocity (RCV). Line calculated using expression (14) from main text and K_m_ = [S_T_] = 10. **C** Hanes plots of eMM rate expressions. Lines drawn using *k_3_* = K_m_ = [E_T_] =1 (dot-dashed black line, expression (27)) or *k_3_* = K_m_ = [S_T_] = 1 (dashed red line, expression (28)). **D** Lineweaver-Burk plots of eMM rate expressions. Lines drawn using *k_3_* = K_m_ = [E_T_] = 1 (dot-dashed black line, expression (25)) or *k_3_* = K_m_ = [S_T_] =1 (dashed red line, expression (26)).

### Reversible monosubstrate reactions

Most chemical reactions in a cell are reversible. The CF approach, assuming FE conditions and two intermediate enzyme complexes ([ES] and [EP]), can be used to calculate the velocity of a reversible enzyme-catalysed reaction in the presence of both substrate and product (Table 1, Supplemental Material 1D). The expression derived is similar to the one derived assuming [E_T_] ≪ [S_T_]. Further, if [E_T_] ≈ 0, the eMM expression can be reduced to the sMM one. However, if expression in Table 1 is rearranged to obtain a single denominator, the additional term in [E_T_] becomes very complex:

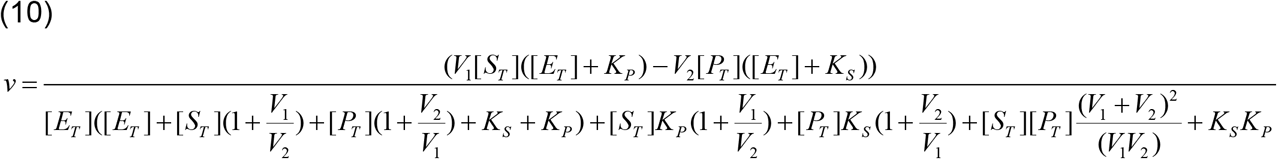

This tendency to complexity is shared with other approaches. Thus, in the hands of this researcher, assuming other conditions, such as QSS, leads to expressions that depart from the simplicity scope of this work (not shown). Nevertheless, those may still describe the process accurately.

### Integrated form of the QSS, eMM expression for irreversible monosubstrate reactions

Expression (9) is amenable to integration and, as expected, it also provides an expression similar to that obtained using standard assumptions (Table 1 and Supplementary Material 1E):

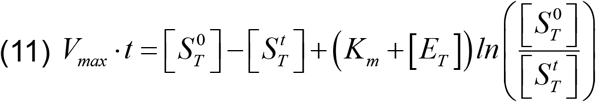

Needless to say that, since there is necessarily no changes in the total amount of enzyme over time, no expression for the integrated form of (9) using [E_T_] as a variable is meaningful.

### Reactions at constant [S_T_]

Similar to the iMM, velocity estimations by eMM at constant [S_T_] shows a hyperbolic behaviour with respect to the amount of [E_T_] in the system (Fig. 2A). Therefore, under those conditions, there exists a maximum asymptotic catalytic velocity attainable that can be defined as

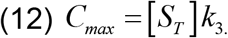

Similarly, we could define as Relative Catalytic Velocity (RCV) the *ad-hoc* parameter

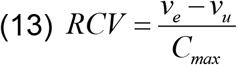

Where *v_e_* is the experimentally observed enzyme-catalysed velocity, *v_u_* is the velocity observed in the absence of enzyme under the same conditions, and *C_max_* the maximum catalysed velocity attainable at the set [S_T_]. Under most circumstances, *vu* is several orders of magnitude smaller than both *v_e_* and *C_max_*; therefore, RCV may be safely approximated by the ratio *v_e_/C_max_*. Hence, RCV should obey:

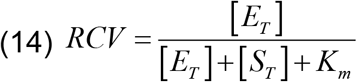

From this, it is possible to demonstrate that, at constant concentration of [S_T_], the concentration of enzyme needed to attain a certain proportion of *C_max_* (RCV) would be:

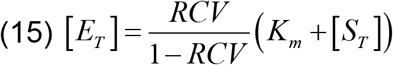

This expression shows that, while RCV is small, the amount of [E_T_] necessary to attain it increases in a near-linear fashion ([*E_T_*] ≈ *RCV*(*K_m_* + [*S_T_*]), but as the values of RCV progress, the values for [E_T_] increase exponentially (Figure 2B).

An example for the use of these expressions could be yeast soluble pyrophosphatase isoform 1 (Ipp1p). This is a well characterised enzyme that hydrolyses Mg•pyrophosphate into two orthophosphate molecules in a *bona fide* irreversible reaction. The K_m_ of that enzyme is typically estimated in the micromolar range, 8 μM being reported earlier [11]. The concentration of the enzyme can be estimated as ~2 μM from the values reported for its median abundance (42812 molecules/cell) and average cell volume (42 μm^3^) [12]. The pyrophosphate concentration in the cell is not precisely known, but probably ranges between 1 and 100 μM [13], with 10 μM being a safe option. Assuming these conditions to be representative of the cell ones, Ipp1p would be working at RCV ≈ 0.1. Similarly, figures ranging from 0.001 (Pyk2p) to 0.3 (Fba1p) can easily be obtained for many of the enzymes involved in yeast glycolysis [14].

### Linearisations of eMM expressions

Although largely superseded by non-linear regression for parameter estimation, linearisations of the rate equations are still useful to show graphically differences between cases or reaction conditions. The CF and QSS-derived rate law (9), and its analogous obtained under FE assumption (Table 1), can also be linearised provided either [E_T_] or [S_T_] are kept constant (Table 1 and Figure 2C and 2D). Among these linearisations are the most commonly ones, namely Lineweaver-Burk and Hanes linearisations. Also, similar to those linearisations, the slopes and intersection points are symmetrical, i.e., the intersection point from a Lineweaver-Burk plot is equivalent to the slope from a Hanes plot and vice-versa.

### Inhibition and activation expressions

CFs were also used to derive expressions for the most common linear inhibition mechanisms under QSS assumptions (Supplementary Material 2). However, in this case, it was also assumed that [I] ≫ [E_T_] ≈ [S_T_], which means that [I_T_] ≈ [I]. This assumption simplifies the expressions and may well be a real situation in the cell.

In all cases, the expressions derived were similar to those obtained under the sMM assumption of negligible concentration of enzyme (Table 2). Noticeably, the enzyme concentration in the denominator also formed a term with the concentration of inhibitor and its equilibrium constant in the case of allosteric inhibition mechanisms (uncompetitive and mixed uncompetitive-competitive). When [ES]_a_ was plotted versus the whole range of [S_T_] and [E_T_], it was observed that a competitive inhibitor at the same conditions of concentration and binding constant values, was much less effective than inhibitors with mechanisms that implied allosteric interactions (Figure 3A). This was also observed in plots with sMM expressions but it was less dramatic (data not shown). Further, using eMM expressions, a comparison between the expected behaviour of those types of inhibitors at [E_T_] ≫ K_m_ and [E_T_] ≈ K_m_ revealed that, in the case of [E_T_] ≫ K_m_, at [S_T_] up to *ca 2x*K_m_, competitive inhibitors are more effective than uncompetitive inhibitors. This was not surprising since sMM expressions already revealed that behaviour (data not shown). However, at [E_T_] ≈ K_m_, uncompetitive inhibitors were more effective than competitive inhibitors in the whole [S_T_] range (Figure 2B and 2C). On the other hand, mixed competitive-uncompetitive inhibitors showed an intermediate situation and were consistently predicted more effective than competitive inhibitors at any concentrations of [E_T_] and [S_T_], in agreement with known behaviour revealed by sMM expressions (data not shown).

**Figure 3.**
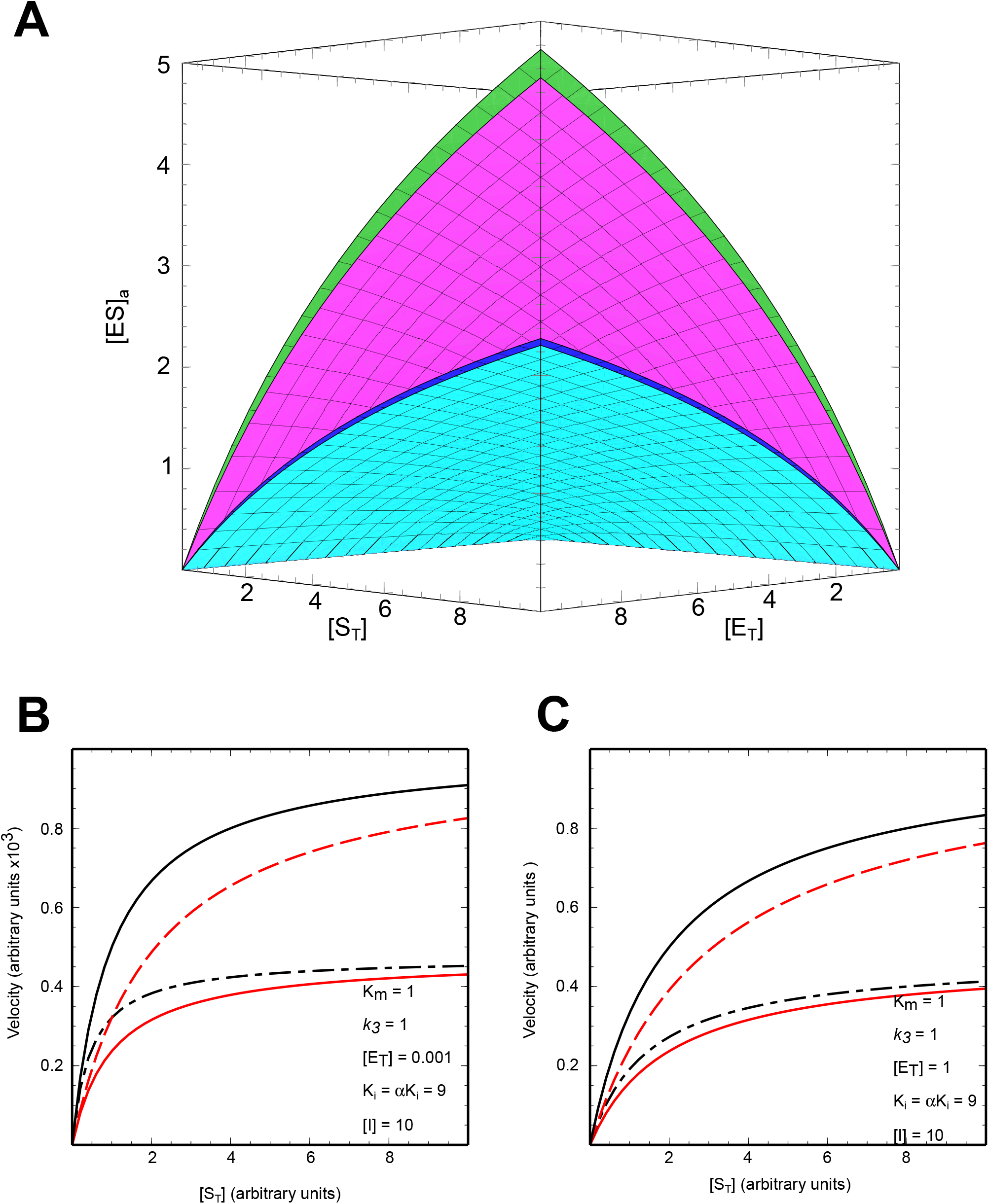
Behaviour of enzyme inhibition expressions obtained using CF approach. **A** Approximated [ES] ([ES]_a_) over [S_T_] and [E_T_]. [ES]_a_ obtained using extended Michaelis-Menten (eMM; expression (7); green surface), competitive inhibition (expression (32); magenta surface), uncompetitive inhibition (expression (34); blue surface) and mixed competitive-uncompetitive/non-competitive inhibition (expressions (36) and (38); cyan surface). **B** Velocity estimated from the different types of inhibition at a constant low concentration of [E_T_] = 0.001 *x* K_m_. Solid black line, no inhibition (expression (9)); dashed red line, competitive inhibition (expression (32)); dot-dashed black line, uncompetitive inhibition (expression (34)); solid red line, mixed/non-competitive inhibition (expressions (36) and (38)). **C** Velocity estimated from the different types of inhibition at [E_T_] = K_m_. Solid black line, no inhibition (expression (9)); dashed red line, competitive inhibition (expression (32)); dot-dashed black line, uncompetitive inhibition (expression (34)); solid red line, mixed/non-competitive inhibition (expressions (36) and (38)). All surfaces and lines calculated using *k_3_* = K_m_ = 1, [I] = 10 and K_i_ = αK_i_ = 9.

### Non-linear inhibition, activation and linearisation of eMM inhibition expressions

Similar to the previous eMM equations and to those obtained under sMM assumptions, eMM equations describing inhibition can be linearised to provide Lineweaver-Burk or Hanes plots (Figure 4). Further, similar to the non-inhibition situation, the values for the slopes and intercepts look transposed when comparing those two types of linearisation.

**Figure 4.**
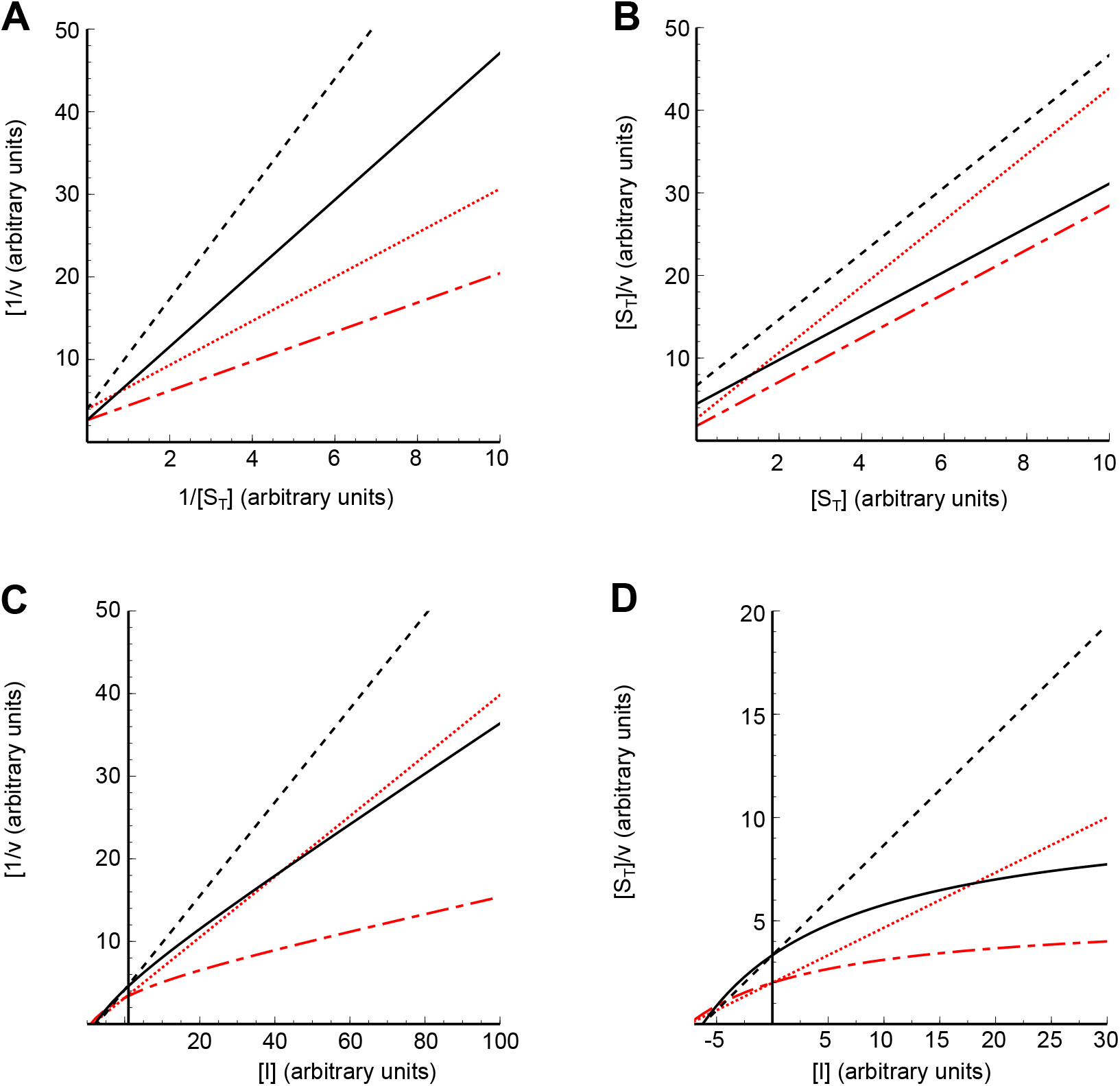
Plots of eMM and sMM linearised equations for total (linear) and partial (non-linear) inhibition mechanisms. All plots correspond to mixed uncompetitive-competitive inhibition as a general representative of all mechanisms. Linear inhibition corresponds to expressions (35) (sMM) and (36) (eMM); non-linear inhibition corresponds to expressions (39) (sMM) and (40) (eMM); values of parameters for all lines as follows: K_m_ = [E_T_] = 1, *k_3_*=0.75, K_i_=10, αK_i_ = 5 and β = 0 (linear) or β = 0.25 (non-linear). **A** Lineweaver-Burk plots. Solid black line, eMM non-linear inhibition; dotted black line, eMM linear inhibition; dash-dotted red line, sMM non-linear inhibition; dotted red line, sMM linear inhibition. **B** Hanes plots. Lines as in A. **C** Dixon plots, Lines as in A. **D** Cornish-Bowden plots. Lines as in A.

Furthermore, the CF approach can be extended to the nonlinear types of inhibition or activation (Table 2). In the case of nonlinear small molecule-enzyme interactions, Lineweaver-Burk and Hanes linearisations provide straight lines similar to those obtained with sMM expressions. Moreover, when Dixon (DX) or Cornish-Bowden (CB) plots are used, bent lines are produced, in agreement with what is observed using sMM expressions.

Being probably the most informative, these types of plots were also checked for consistency with classical expressions in the case of linear inhibition. CB plots from eMM expressions produced straight lines that, in the case of allosteric inhibitors, intersected in the second or third quadrant or on the negative side of the abscissa (depending on the value of α); on the contrary, in the case of competitive inhibitors, parallel lines were observed (Figure 5). This is similar to what is described for sMM expressions [15]. The absolute value of the abscissa at the intersection point of the lines was estimated to be equal to the allosteric equilibrium constant for the binding of the inhibitor to the enzyme (Figure 5 and Table 3). All this is identical to what is found using sMM expressions. However, important differences were observed in Dixon plots. Intersection of the lines did not follow the pattern expected from sMM expressions. Thus, uncompetitive inhibition equations showed convergent lines on DX plots that crossed in the third quadrant (Figure 5B) while parallel lines are typical from sMM equations. In appearance, competitive inhibition, mixed uncompetitive-competitive inhibition and non-competitive inhibition expressions produce DX plots that agree with their sMM counterparts. However, the abscissa absolute values for the line intersection points are all dependent on the values of K_m_ and [E_T_] in the system (Table 3). Only in the case of non-competitive inhibition, the abscissa absolute value corresponds to K_i_, as it is the case of sMM expressions.

**Figure 5.**
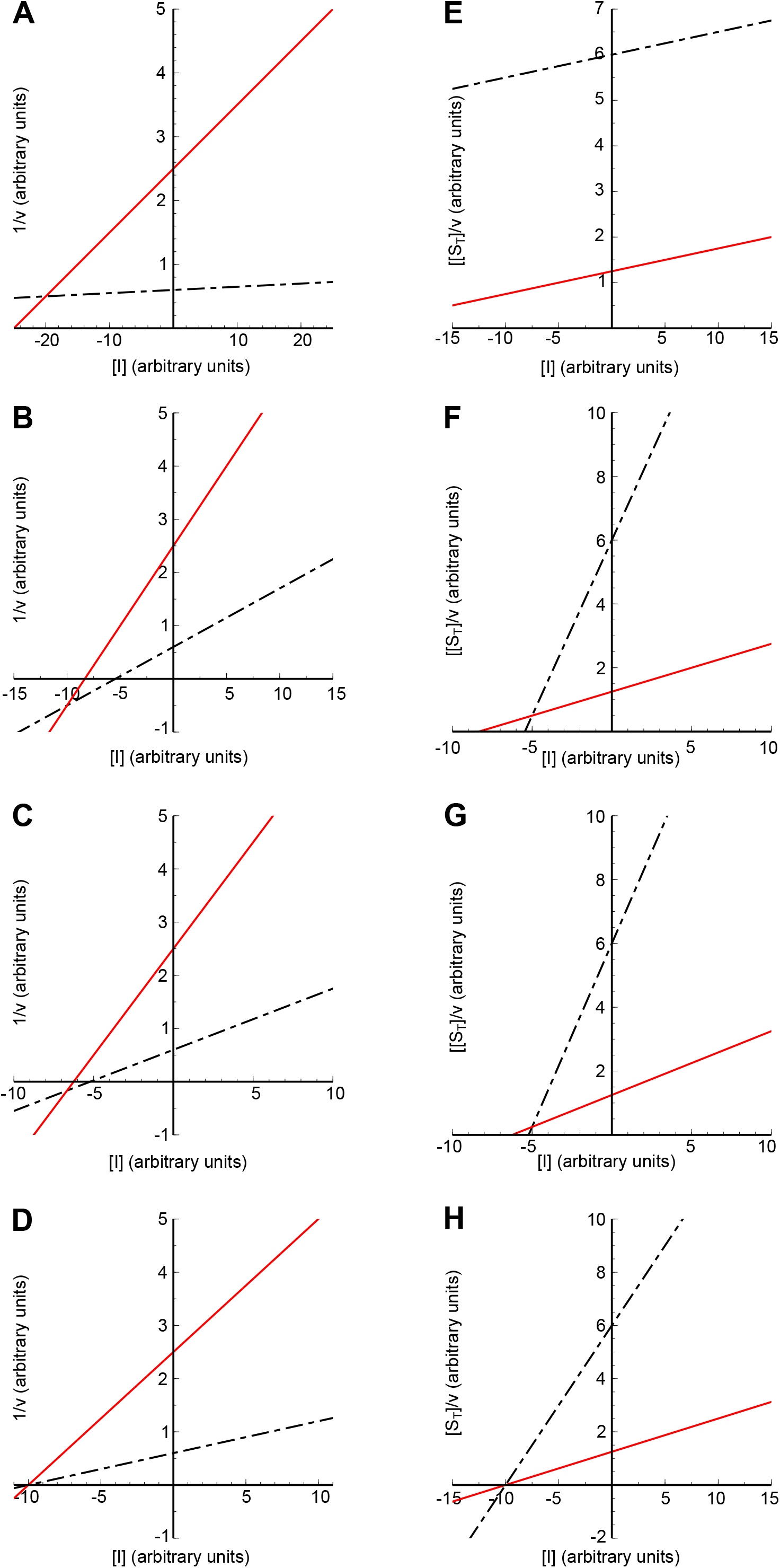
Dixon and Cornish-Bowden plots for eMM linear inhibition mechanisms. All lines drawn using equations (32) (competitive), (34) (uncompetitive), (36) (mixed competitive-uncompetitive) and (38) (non-competitive) with the following parameters: K_m_ = [E_T_] = 1, *k_3_* = 2, K_i_ = 10, αK_i_ = 5. Solid red lines, [S_T_] = 0.5; dash-dotted black lines [S_T_] = 10. Panels **A** and **E**, competitive inhibition; panels **B** and **F**, uncompetitive inhibition, panels **C** and **G**, mixed competitive-uncompetitive inhibition; panels **D** and **H**, non-competitive inhibition. Panels **A** to **D**, Dixon plots; panels **E** to **H**, Cornish-Bowden plots.

## DISCUSSION

Derivation of expressions predicting the velocity of monosubstrate enzyme-catalysed reactions was found possible using a simple algebraic approach. Further, the expressions were similar to those already in use (sMM) and could be reduced to those under the assumption of negligible concentration of [ET]. This helps them to be understood as easy to use extensions of the classical expressions. Nevertheless, these equations only provide approximations to the real value, and, therefore, care should be taken that assumptions are valid when using them. In any case, the expression proposed to estimate the concentration of the ES complex in a monosubstrate irreversible reaction (7) is identical to those proposed earlier [4,5,16,17] and was found to approximate accurately the true value of [ES] from (2) provided (*K_m_* + [*E*] + [*S_T_*]) ≫ [*ES*]. It is true that expressions (7) and (9) are not new but, to the best of this researcher’s knowledge, this is the first report were the continued fraction approach was used to obtain it. This is a simple approach that is capable of different degrees of accuracy, if needed. Also, its simplicity may help students to understand how expressions are derived and experimental researchers to get familiar with them. Furthermore, a wide range of expressions are made available and not only an equation describing the simplest monosubstrate, irreversible case. That should open the possibility for their use in a wide variety of conditions and studies.

Some previous proposals were based on macroscopic parameters different from those experimental biochemists are used to obtain [18]. While the estimation of different parameters may just be a question of use and learning, undoubtedly the possibility of using the same approaches and parameters to extrapolate enzyme behaviour in conditions difficult to mimic in the test tube may be considered an advantage.

The introduction of [E_T_] as a reactant in the denominator of the equations in this study leads to the concept of enzyme saturation in a reaction. This means that increasing the concentration of enzyme in a system above a certain amount may not result in meaningful increases in velocity. Therefore, equation (14) can be a useful tool to understand the cellular effort needed to modulate cell concentrations of metabolites, the feasibility in each context of gene regulation of protein expression or to evaluate the investment needed to attain an acceptable velocity in a biotechnological setting.

Linearisations are used extensively by biochemists to show results and point to differences in enzyme action. The expressions presented in this work can also be linearised using, at least, some of the common approaches in use for sMM. They provide lines that, in most cases, behave just like the sMM ones. However, as other linearisations are tested, exceptions, like those observed in Dixon plots, may appear.

One particularity of these linearisations for a simple irreversible reaction is that, at least theoretically, it could be possible to estimate the value of *k_3_* (or *k_cat_*) in non-pure samples, such as extracts. That determination would need sufficient different amounts of extract assayed for activity maintaining a constant [S_T_]. The velocities obtained could be used in a plot of [E_T_]/v *vs* [E_T_], where [E_T_] can be expressed as fold over the sample having the smallest amount of extract. The reciprocal of the slope of that straight line would equal [S_T_]*k_3_*. Since [S_T_] is a known parameter in the assay, *k_3_* could be estimated. Nevertheless, maintaining the QSS assumption valid in those experiments may well be the major hurdle. Optimally, to ensure accuracy of the estimation, several of the determinations should need to fall in the non-linear part of the curve. In other words, those determinations should need to be made in conditions where [S_T_] is close to that of [E_T_] and, thus, it can be very difficult to assume [P] ≈0 or that variations in [ES] are negligible during the assay. In any case, in special cases, it might be a possibility.

Although similar to the classical expressions, the equations presented here already revealed some differences that may be important to understand enzyme and inhibitor action *in vivo*. The value of K_m_ in sMM it is often described as (and sometimes used as a definition) the concentration of substrate providing 1/2V_max_. On the other hand, the catalytic efficiency (the slope of the line describing first order kinetics), is a parameter often used to compare catalysis on different substrates and even isoenzyme performance. However, eMM equations showed that, in a general setting, [E_T_] needs to be taken into account in those cases. While the issue of the K_m_ may not have important consequences in experimental biochemistry and the values of the catalytic efficiencies at [E_T_] = 0 could still be used to compare different catalytic situations, other differences may be more relevant. For example, the increase in inhibitory power of uncompetitive inhibitors that is only observed at [E_T_] ≈ [S_T_] ≈ K_m_. That difference, together with the general differences in effect between allosteric and catalytic inhibitors predicted by these expressions, may be useful to understand inhibitor action *in* vivo or to direct the search for new drugs in a more selective way.

Finally, these expressions cover only a small part of the situations that can be faced in relation to enzyme kinetics. Among the most important ones not covered by this work are multisubstrate reactions and allosteric or cooperative enzymes. Efforts are in place now to deduce equations able to attend to those cases. On the whole, the expressions proposed here may be useful tools for the translation *in vivo* of *in vitro* experimental data or for modelling *in vivo* and biotechnological processes.

## Acknowledgements

The author thanks Ms E. A. de Magalhães for her help in this study. This work is dedicated to late Drs D. T. Cooke and D. T. Clarkson for lifelong teaching through example.

The author declares not to have received funds for the development of this study and no conflicts of interest.

## Supporting Information

### Supplementary Materials 1

Derivation of the equations for the simple cases: The exact solution for irreversible QSS monosubstrate reaction, single monosubstrate irreversible FE eMM equation, single monosubstrate irreversible QSS eMM equation, reversible single monosubstrate FE eMM equation, and integrated QSS eMM equation.

### Supplementary Materials 2

Derivation of the eMM equations for inhibition and activation under QSS. Competitive linear inhibition, uncompetitive linear inhibition, mixed competitive-uncompetitive linear inhibition, non-competitive linear inhibition, and general inhibition/activation model.

**Table 1.** Comparison of velocity expressions for monosubstrate enzyme-catalysed reactions.

|  | sMM | eMM | Linearisations (eMM) |
| --- | --- | --- | --- |
| FE<br>(Irr.) | $v = \frac{V_{max} \cdot [S_T]}{[S_T] + K_S} \quad (16)$ | $v = \frac{V_{max} \cdot [S_T]}{[E_T] + [S_T] + K_S} \quad (9)$ | $\frac{1}{v} = \frac{1}{[S_T]} \cdot \frac{[E_T] + K_S}{V_{max}} + \frac{1}{V_{max}} \quad (17)$ |
| | | | $\frac{1}{v} = \frac{1}{[E_T]} \cdot \frac{[S_T] + K_S}{C_{max}} + \frac{1}{C_{max}} \quad (18)$ |
| | | | $\frac{[S_T]}{v} = [S_T] \frac{1}{V_{max}} + \frac{[E_T] + K_S}{V_{max}} \quad (19)$ |
| | | | $\frac{[E_T]}{v} = [E_T] \frac{1}{C_{max}} + \frac{[S_T] + K_S}{C_{max}} \quad (20)$ |
| FE<br>(Rev.) | $v = \frac{V_1[S]}{[S] \left(1 + \frac{V_1}{V_2}\right) + K_S} - \frac{V_2[P]}{[P] \left(1 + \frac{V_2}{V_1}\right) + K_P} \quad (21)$ | $v = \frac{V_1[S_T]}{[E_T] + [S_T] \left(1 + \frac{V_1}{V_2}\right) + K_S} - \frac{V_2[P_T]}{[E_T] + [P_T] \left(1 + \frac{V_2}{V_1}\right) + K_P} \quad (22)$ | |
| QSS<br>(Irr.) | $v = \frac{V_{max} \cdot [S_T]}{[S_T] + K_m} \quad (23)$ | $v = \frac{V_{max} \cdot [S_T]}{[E_T] + [S_T] + K_m} \quad (24)$ | $\frac{1}{v} = \frac{1}{[S_T]} \cdot \frac{[E_T] + K_m}{V_{max}} + \frac{1}{V_{max}} \quad (25)$ |
| | | | $\frac{1}{v} = \frac{1}{[E_T]} \cdot \frac{[S_T] + K_m}{C_{max}} + \frac{1}{C_{max}} \quad (26)$ |
| | | | $\frac{[S_T]}{v} = [S_T] \frac{1}{V_{max}} + \frac{[E_T] + K_m}{V_{max}} \quad (27)$ |
| | | | $\frac{[E_T]}{v} = [E_T] \frac{1}{C_{max}} + \frac{[S_T] + K_m}{C_{max}} \quad (28)$ |
| QSS<br>(Irr.) | $V_{max} \cdot t = [S_T^0] - [S_T^t] + K_m \ln \left( \frac{[S_T^0]}{[S_T^t]} \right) \quad (29)$ | $V_{max} \cdot t = [S_T^0] - [S_T^t] + (K_m + [E_T]) \ln \left( \frac{[S_T^0]}{[S_T^t]} \right) \quad (11)$ | $\frac{([S_T^0] - [S_T^t])}{t} = V_{max} - (K_m + [E_T]) \ln \left( \frac{[S_T^0]}{[S_T^t]} \right) \left( \frac{1}{t} \right) \quad (30)$ |

FE: Fast Equilibrium assumption, QSS: Quasi-steady state assumption, Irr.: Irreversible reaction, Rev.: Reversible reaction, sMM: standard Michaelis-Menten, eMM: extended Michaelis-Menten

**Table 2.**
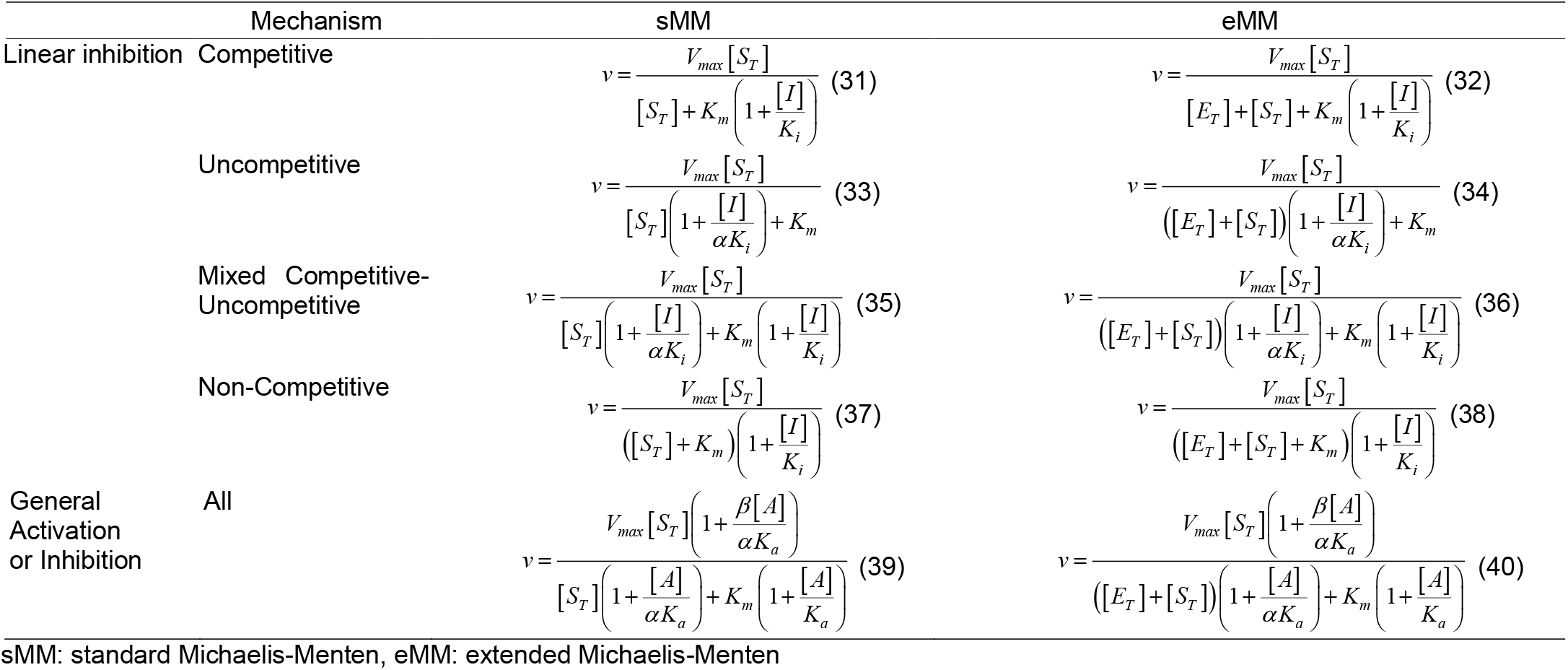
Comparison of velocity expressions for monosubstrate, irreversible, enzyme-catalysed reactions affected by a non-covalent modifier (inhibitor or activator). All expressions derived assuming QSS.

**Table 3.** Abscissa values for the intersection points of lines created using eMM linear inhibition expressions in Dixon and Cornish-Bowden plots.

| | Dixon ($1/v$ vs $[I]$) | Cornish-Bowden ($[S_T]/v$ vs $[I]$) |
| --- | --- | --- |
| Competitive | $-K_i \frac{[E_T] + K_m}{K_m}$ | No intersection |
| Uncompetitive | $-\alpha K_i \frac{[E_T] + K_m}{[E_T]}$ | $-\alpha K_i$ |
| Mixed Competitive-Uncompetitive | $-\alpha K_i \frac{[E_T] + K_m}{[E_T] + \alpha K_m}$ | $-\alpha K_i$ |
| Non-Competitive | $-K_i$ | $-K_i$ |

