## Supplemental Material 1 for "Mathematical expressions describing enzyme velocity and inhibition at high enzyme concentration"

**Derivation of enzyme kinetics equations under conditions of high concentration of enzyme. Simple cases.**

**(A) Derivation of the exact solution for an irreversible reaction under quasi-steady state conditions (QSS).**

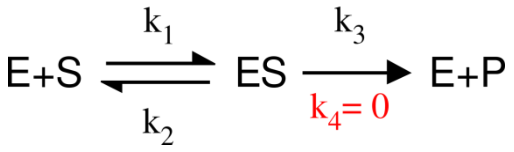


The assumption of the irreversibility of the reaction is a consequence of focusing on the initial states of the reaction (t=0). Under that condition, [P] ≈ 0, which leads to [E][P]$k_4$ = 0. Despite being erroneous, for simplicity sake only, that is depicted in the scheme above and in the following ones as $k_4$ = 0.

Under these conditions and assumptions, a single substrate irreversible reaction will be described by the following expressions:

(1) $[E_T] = [E] + [ES]$

(2) $[S_T] = [S] + [ES]$

(3) $\frac{d[ES]}{dt} = 0 = k_1[E]\cdot[S] - k_2[ES] - k_3[ES]$

(4) $\frac{d[P]}{dt} = k_3[ES]$

(5) $K_m = \frac{k_2 + k_3}{k_1}$

From (1), (2), (3) we can deduce:

(6) $0 = ([E_T]-[ES])([S_T]-[ES])k_1 - [ES]k_2 - [ES]k_3$

After expanding the product, gathering terms in [ES] and substituting (5):

(7) $0 = [ES]^2 - [ES]\cdot(K_m + [E_T] + [S_T]) + [S_T]\cdot[E_T]$

The exact solution for this equation is:

(8) $[ES] = \frac{1}{2}\left((K_m + [S_T] + [E_T]) \pm \sqrt{(K_m + [S_T] + [E_T])^2 - 4\cdot[S_T]\cdot[E_T]}\right)$

The positive sign solution of (8) makes no chemical sense, exemplified by the case that in of absence of substrate: $[ES] = K_m + [E_T]$

Hence, the rate law is:

(9) $\frac{d[P]}{dt} = \frac{k_3}{2}\left((K_m + [S_T] + [E_T]) - \sqrt{(K_m + [S_T] + [E_T])^2 - 4\cdot[S_T]\cdot[E_T]}\right)$

**(B) Derivation of the first and second degree approximants using continuous fractions (eMM) for a single substrate irreversible reaction under QSS assumptions.**

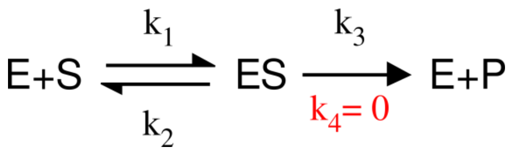


As in (A) above:

(1) $[E_T]=[E]+[ES]$

(2) $[S_T]=[S]+[ES]$

(3) $\dfrac{d[ES]}{dt}=0=k_1[E]\cdot[S]-k_2[ES]-k_3[ES]$

(4) $\dfrac{d[P]}{dt}=k_3[ES]$

(5) $K_m=\dfrac{k_2+k_3}{k_1}$

(10) $V_{max}=k_3[E_T]$

As in (A), using (1) and (2) into (3), expanding the product, gathering terms in [ES] and substituting (5):

(7) $0=[ES]^2-[ES]\cdot\left(K_m+[E_T]+[S_T]\right)+[S_T]\cdot[E_T]$

Which can be written also as:

(11) $0=[ES]\cdot\left([ES]-\left(K_m+[E_T]+[S_T]\right)\right)+[S_T]\cdot[E_T]$

Rearranging:

(12) $-[S_T][E_T]=[ES]\cdot\left([ES]-\left(K_m+[E_T]+[S_T]\right)\right)$

Using the continued fraction approximation approach we can obtain both solutions for [ES]. The positive sign solution would be obtained by dividing both sides of equation (12) by [ES] and later substracting the independent term on the right hand side:

(13) Positive sign solution: $[ES]=\left(K_m+[E_T]+[S_T]\right)-\dfrac{[S_T]\cdot[E_T]}{[ES]}$

However, this solution will be discarded for the reasons stated for (8) above.

For the negative sign solution, both sides of equation (12) are divided by the term in the parenthesis on the right hand side.

(14) Negative sign solution: $[ES]=\dfrac{[S_T]\cdot[E_T]}{\left(K_m+[E_T]+[S_T]\right)-[ES]}$

Assuming $\left(K_m+[E_T]+[S_T]\right)\gg[ES]$, the term [ES] in the denominator can be neglected and equation (14) can be approximated to:

(15) $[ES]\simeq\frac{[S_T]\cdot[E_T]}{K_m+[E_T]+[S_T]}$

From (15) and (4), we finally obtain (16) $\frac{d[P]}{dt}=\frac{[S_T]\cdot[E_T]\cdot k_3}{K_m+[E_T]+[S_T]}$

Using (6), expression (16) can also be writen as (17) $\frac{d[P]}{dt}=\frac{V_{max}\cdot[S_T]}{K_m+[E_T]+[S_T]}$

Higher degree approximations can be obtained from (14) by recursively subtituiting [ES] in the denominator with the right hand side of the equation (14) again and neglecting [ES] in the last iterative substitution. Thus, a **second order approximant** would be:

(18) $[ES]=\frac{[S_T]\cdot[E_T]}{\left(K_m+[E_T]+[S_T]\right)-\frac{[S_T]\cdot[E_T]}{\left(K_m+[E_T]+[S_T]\right)}}$

Rearranging:

(19) $[ES]=\frac{\left(K_m+[E_T]+[S_T]\right)[S_T][E_T]}{\left(K_m+[E_T]+[S_T]\right)^2-[S_T][E_T]}$

**(C) Derivation of the first degree approximant using continuous fractions (eMM) for a single substrate irreversible reaction under fast equilibrium (FE) assumptions.**

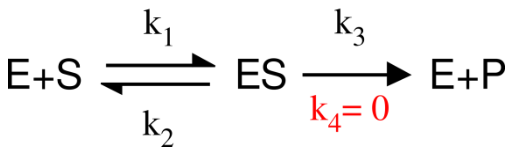


The expressions describing this reaction are:

(1) $[E_T]=[E]+[ES]$

(2) $[S_T]=[S]+[ES]$

(4) $\frac{d[P]}{dt}=k_3[ES]$

(10) $V_{max}=k_3[E_T]$

(20) $K_S=\frac{k_2}{k_1}=\frac{[E]\cdot[S]}{[ES]}$

We can rearrange (20):

(21) $0=[E]\cdot[S]-\frac{k_2}{k_1}[ES]$

As in (B) above, we can substitute (1) and (2) into (21), expand the product, rearrange to gather terms in [ES] and substitute microscopic rate constants by the binding equilibrium constant:

(22) $0=[ES]^2-[ES]\cdot(K_s+[E_T]+[S_T])+[S_T]\cdot[E_T]$

Using the continued fraction approach as above, we can reach:

(23) $[ES]=\frac{[S_T]\cdot[E_T]}{(K_s+[E_T]+[S_T])-[ES]}$

Assuming $(K_s+[E_T]+[S_T])\gg[ES]$

(24) $[ES]\simeq\frac{[S_T]\cdot[E_T]}{K_s+[E_T]+[S_T]}$

Finally, the rate law would be (25) $\frac{d[P]}{dt}=\frac{[S_T]\cdot[E_T]\cdot k_3}{K_s+[E_T]+[S_T]}$

Which can also be writen as (26) $\frac{d[P]}{dt}=\frac{V_{max}\cdot[S_T]}{K_s+[E_T]+[S_T]}$

**(D) Derivation of the first degree approximant using continuous fractions (eMM) for a single substrate reversible reaction with two enzyme complexes under fast equilibrium (FE) assumptions.**

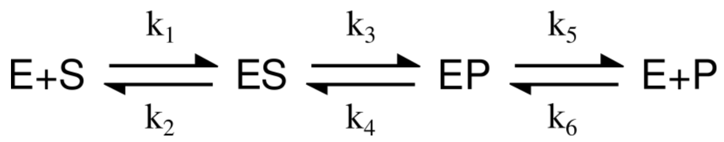


The expressions describing this reaction are:

(27) $[E_T]=[E]+[ES]+[EP]$

(2) $[S_T]=[S]+[ES]$

(28) $[P_T]=[P]+[EP]$

(29) $v=\frac{d[P]}{dt}=k_3[ES]-k_4[EP]$

(10) $V_{max}^{S\to P}=V_1=k_3[E_T]$

(30) $V_{max}^{P\to S}=V_2=k_4[E_T]$

(20) $K_S=\frac{k_2}{k_1}=\frac{[E]\cdot[S]}{[ES]}$

(31) $K_P=\frac{k_5}{k_6}=\frac{[E]\cdot[P]}{[EP]}$

(32) $K_C=\frac{k_4}{k_3}=\frac{V_2}{V_1}=\frac{[ES]}{[EP]}$

We can rearrange the expression (32) to obtain:

(33) $[EP]\frac{V_2}{V_1}=[ES]$ and also (34) $[ES]\frac{V_1}{V_2}=[EP]$

Substituting either (33) or (34) into the enzyme mass conservation expression (27) we can obtain two different expressions:

(35) $[E]=[E_T]-[ES]\left(1+\frac{V_1}{V_2}\right)$ and (36) $[E]=[E_T]-[EP]\left(1+\frac{V_2}{V_1}\right)$, respectively.

In the substrate-enzyme binding equilibrium (20), we can substitute [E] with (35) and [S] with $[S_T]$-[ES]. After rearranging:

(37) $[ES]K_S=\left([E_T]-[ES]\left(1+\frac{V_1}{V_2}\right)\right)([S_T]-[ES])$

Expanding the product and leaving zero at the right hand side

(38) $$0=[ES]^2\left(1+\frac{V_1}{V_2}\right)-[ES]\left([E_T]+[S_T]\left(1+\frac{V_1}{V_2}\right)+K_S\right)+[E_T][S_T]$$

Eliminate the coeficient in the squared term:

(39) $$0=[ES]^2-[ES]\frac{1}{\left(1+\frac{V_1}{V_2}\right)}\left([E_T]+[S_T]\left(1+\frac{V_1}{V_2}\right)+K_S\right)+\frac{1}{\left(1+\frac{V_1}{V_2}\right)}[E_T][S_T]$$

The continued fraction will be:

(40) $$[ES]=\frac{\frac{1}{\left(1+\frac{V_1}{V_2}\right)}[E_T][S_T]}{\frac{1}{\left(1+\frac{V_1}{V_2}\right)}\left([E_T]+[S_T]\left(1+\frac{V_1}{V_2}\right)+K_S\right)-[ES]}$$

Simplified for a first degree approximant:

(41) $$[ES]\simeq\frac{[E_T][S_T]}{[E_T]+[S_T]\left(1+\frac{V_1}{V_2}\right)+K_S}$$

A similar procedure using the product-enzyme binding equilibrium (31) and expression (36) will render:

(42) $$[EP]\simeq\frac{[E_T][P_T]}{[E_T]+[P_T]\left(1+\frac{V_2}{V_1}\right)+K_P}$$

The velocity expression is then:

(43) $$v=\frac{V_1[S_T]}{[E_T]+[S_T]\left(1+\frac{V_1}{V_2}\right)+K_S}-\frac{V_2[P_T]}{[E_T]+[P_T]\left(1+\frac{V_2}{V_1}\right)+K_P}$$

This is a simple enough expression. However, if a single denominator is sought out, then:

(44) $$v=\frac{V_1[S_T]\left([E_T]+[P_T]\left(1+\frac{V_2}{V_1}\right)+K_P\right)-V_2[P_T]\left([E_T]+[S_T]\left(1+\frac{V_1}{V_2}\right)+K_S\right)}{\left([E_T]+[S_T]\left(1+\frac{V_1}{V_2}\right)+K_S\right)\left([E_T]+[P_T]\left(1+\frac{V_2}{V_1}\right)+K_P\right)}$$

Upon expansion, cancelation of terms and rearranging we obtain the final expression (45):

$$\frac{V_1[S_T]]([E_T]+K_P)-V_2[P_T]([E_T]+K_S)}{[E_T]\left([E_T]+[S_T]\left(1+\frac{V_1}{V_2}\right)+[P_T]\left(1+\frac{V_2}{V_1}\right)+K_S+K_P\right)+[S_T]K_P\left(1+\frac{V_1}{V_2}\right)+[P_T]K_S\left(1+\frac{V_2}{V_1}\right)+[S_T][P_T]\frac{(V_1+V_2)^2}{V_1V_2}+K_SK_P}$$

Which is similar to the expression obtained under standard assumptions of [S] >> [$E_T$] and that it can be reduced to that expression if [$E_T$] ≈ 0:

$$v = \frac{V_1[S]}{[S]\left(1+\frac{V_1}{V_2}\right)+K_S} - \frac{V_2[P]}{[P]\left(1+\frac{V_2}{V_1}\right)+K_P} = \frac{V_1[S_T]K_P - V_2[P_T]K_S}{[S_T]K_P\left(1+\frac{V_1}{V_2}\right)+[P_T]K_S\left(1+\frac{V_2}{V_1}\right)+[S_T][P_T]\frac{(V_1+V_2)^2}{V_1V_2}+K_SK_P}$$

**(D) Integrated form of the continued fraction first degree approximant (eMM) for a single substrate irreversible reaction obtained under QSS assumptions.**

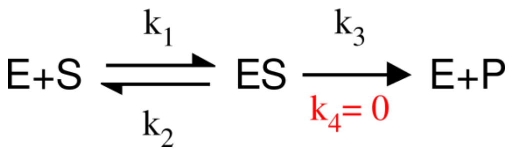


The initial reaction velocity expression obtained was:

(16) $\frac{d[P]}{dt} = \frac{[S_T]\cdot[E_T]\cdot k_3}{K_m + [E_T] + [S_T]}$

Since (46) $\frac{d[P]}{dt} = -\frac{d[S]}{dt}$

Expression (46) can be used in (16) and, after rearranging differentials and taking limits:

(47) $-\int_0^t [E_T]\cdot k_3 dt = \int_{[S_T^0]}^{[S_T^t]} \frac{K_m + [E_T] + [S_T]}{[S_T]} d[S]$

Rearranging the quotient and resolving the first integral

(48): $-[E_T]\cdot k_3 \cdot t = \int_{[S_T^0]}^{[S_T^t]} 1 d[S] + \int_{[S_T^0]}^{[S_T^t]} \left( (K_m + [E_T]) \frac{1}{[S_T]} \right) d[S]$

Finally (49) $[E_T]\cdot k_3 \cdot t = [S_T^0] - [S_T^t] + (K_m + [E_T]) ln \left( \frac{[S_T^0]}{[S_T^t]} \right)$

Expression (49) can also be writen as (50) $V_{max} \cdot t = [S_T^0] - [S_T^t] + (K_m + [E_T]) ln \left( \frac{[S_T^0]}{[S_T^t]} \right)$
