## Supplemental Material 2 for "Mathematical expressions describing enzyme velocity and inhibition at high enzyme concentration"

**Derivation of the rate law equations for a monosubstrate enzyme-catalysed reaction, in the presence of an inhibitor, using first degree approximants from continuous fractions (eMM) and assuming QSS conditions.**

### (A) Competitive Inhibition

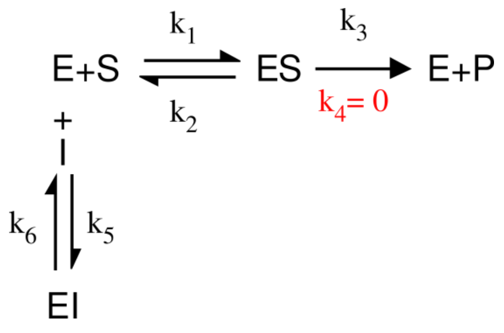


Under the above conditions and assumptions, a single substrate, irreversible reaction will be described by the following expressions:

Assuming [I] >> [$E_T$], i.e. [$I_T$] ≈ [I]

(2) $[S_T] = [S] + [ES]$

(3) $\frac{d[ES]}{dt} = 0 = k_1[E]\cdot[S] - k_2[ES] - k_3[ES]$

(4) $\frac{d[P]}{dt} = k_3[ES]$

(5) $K_m = \frac{k_2 + k_3}{k_1}$

(10) $V_{max} = k_3[E_T]$

(51) $[E_T] = [E] + [ES] + [EI]$

(52) $K_i = \frac{k_6}{k_5} = \frac{[E][I]}{[EI]}$

From (51) and (52) we can deduce:

(53) $[E_T] = [E]\left(1 + \frac{[I]}{K_i}\right) + [ES]$

Dividing both sides of (3) by $k_1$, gathering terms in [ES] and substituting (5), we can express:

(54) $0 = [E][S] - [ES]\cdot K_m$

Substituting (2) and (53) into (54):

(55) $$0=\frac{([E_T]-[ES])}{1+\frac{[I]}{K_i}}\cdot([S_T]-[ES])-[ES]K_m$$

Expanding the product:

(56) $$0=\frac{[E_T][S_T]-[ES][S_T]-[E_T][ES]+[ES]^2}{1+\frac{[I]}{K_i}}-[ES]K_m$$

Multiplying both sides by $1+\frac{[I]}{K_i}$:

(57) $$0=[E_T][S_T]-[ES][S_T]-[E_T][ES]+[ES]^2-[ES]K_m\left(1+\frac{[I]}{K_i}\right)$$

Grouping [ES] terms:

(58) $$0=[ES]^2-[ES]\left([E_T]+[S_T]+K_m\left(1+\frac{[I]}{K_i}\right)\right)+[E_T][S_T]$$

Using the continued fraction approach for the negative sign solution:

(59) $$[ES]=\frac{[E_T][S_T]}{\left([E_T]+[S_T]+K_m\left(1+\frac{[I]}{K_i}\right)\right)-[ES]}$$

Assuming that $\left([E_T]+[S_T]+K_m\left(1+\frac{[I]}{K_i}\right)\right)\gg[ES]$, (59) can be approximated to:

(60) $$[ES]\simeq\frac{[E_T][S_T]}{[E_T]+[S_T]+K_m\left(1+\frac{[I]}{K_i}\right)}$$

Therefore, the rate law is:

(61) $$\frac{d[P]}{dt}=\frac{[E_T][S_T]k_3}{[E_T]+[S_T]+K_m\left(1+\frac{[I]}{K_i}\right)}$$

Which can also be written as:

(62) $$\frac{d[P]}{dt}=\frac{V_{max}[S_T]}{[E_T]+[S_T]+K_m\left(1+\frac{[I]}{K_i}\right)}$$

**(B) Uncompetitive Inhibition**

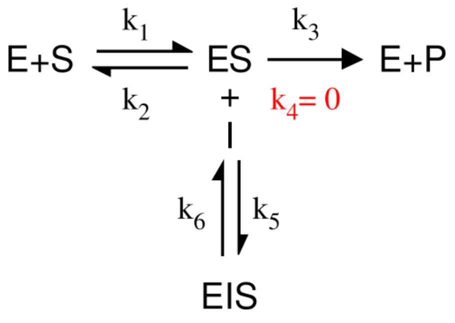


Similarly to the previous case, a single substrate, irreversible reaction will be described by the following expressions:

Assuming [I] >> [$E_T$]

(4) $\frac{d[P]}{dt} = k_3[ES]$

(5) $K_m = \frac{k_2 + k_3}{k_1}$

(10) $V_{max} = k_3[E_T]$

(63) $[E_T] = [E] + [ES] + [ESI]$

(64) $[S_T] = [S] + [ES] + [ESI]$

(65) $\frac{d[ES]}{dt} = 0 = k_1[E] \cdot [S] + k_6[ESI] - k_2[ES] - k_3[ES] - k_5[ES][I]$

(66) $\alpha K_i = \frac{k_6}{k_5} = \frac{[ES][I]}{[ESI]}$

From (66) we get (67) $[ESI] = \frac{[ES][I]k_5}{k_6}$

Substituting (67) into (63) and (64) and grouping terms in [ES]:

(68) $[E_T] = [E] + [ES] \cdot \left(1 + \frac{[I]k_5}{k_6}\right)$ and (69) $[S_T] = [S] + [ES] \cdot \left(1 + \frac{[I]k_5}{k_6}\right)$, respectively.

Substituting (68) and (69) into (65), we obtain: (70)

$$0 = k_1 \cdot \left([E_T] - [ES] \cdot \left(1 + \frac{[I]k_5}{k_6}\right)\right)\left([S_T] + [ES]\left(\frac{[I]k_5}{k_6}\right)\right) + [ES] \cdot \frac{[I]k_5}{k_6} k_6 - k_2[ES] - k_3[ES] - [ES][I]k_5$$

After cancelling the last two terms in $k_5$ and expanding, we obtain: (72)

$$0=[E_T][S_T]k_1-[ES][E_T]k_1\left(1+\frac{[I]k_5}{k_6}\right)-[ES][S_T]k_1\left(1+\frac{[I]k_5}{k_6}\right)+[ES]^2k_1\left(1+\frac{[I]k_5}{k_6}\right)^2-[ES]k_2-[ES]k_3$$

Dividing both sides of the equality by $k_1\left(1+\frac{[I]k_5}{k_6}\right)^2$ and grouping [ES] terms, we obtain:

(73)

$$0=[ES]^2-[ES]\frac{1}{k_1\left(1+\frac{[I]k_5}{k_6}\right)^2}\left([E_T]k_1\left(1+\frac{[I]k_5}{k_6}\right)+[S_T]k_1\left(1+\frac{[I]k_5}{k_6}\right)+k_2+k_3\right)+\frac{1}{k_1\left(1+\frac{[I]k_5}{k_6}\right)^2}[E_T][S_T]k_1$$

Using the continued fraction approach for the negative sign solution:

(74) $$[ES]=\frac{\frac{-1}{k_1\left(1+\frac{[I]k_5}{k_6}\right)^2}[E_T][S_T]k_1}{\frac{-1}{k_1\left(1+\frac{[I]k_5}{k_6}\right)^2}\left([E_T]k_1\left(1+\frac{[I]k_5}{k_6}\right)+[S_T]k_1\left(1+\frac{[I]k_5}{k_6}\right)+k_2+k_3\right)-[ES]}$$

Assuming [ES] << rest of the denominator, we can neglect [ES] and, therefore, also cancel the coefficients $\frac{-1}{k_1\left(1+\frac{[I]k_5}{k_6}\right)^2}$. If we then divide numerator and denominator by $k_1$

(75) $$[ES]\simeq\frac{[E_T][S_T]}{[E_T]\left(1+\frac{[I]k_5}{k_6}\right)+[S_T]\left(1+\frac{[I]k_5}{k_6}\right)+\frac{k_2+k_3}{k_1}}$$

Substituting (75) into (4), grouping terms and attending the definition of the macroscopic kinetic constants (5), (10) and (66), we finally obtain the rate law:

(76) $$\frac{d[P]}{dt}=\frac{V_{max}[S_T]}{\left([E_T]+[S_T]\right)\left(1+\frac{[I]}{\alpha K_i}\right)+K_m}$$

**(C) Mixed Competitive-Uncompetitive Inhibition:**

For a Non-Competitive Inhibition, the same derivation applies. However, $k_6/k_5 = k_8/k_7$, *and therefore* $K_i = \alpha K_i$

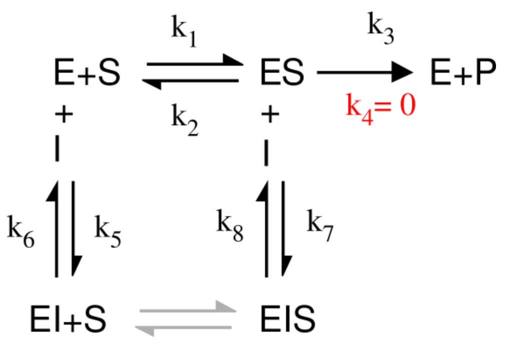


Similarly to the previous cases, a single substrate, irreversible reaction, in the presence of a mixed-type inhibitor will be described by the following expressions:

Assuming [I] >> [$E_T$]

(4) $\frac{d[P]}{dt} = k_3[ES]$

(5) $K_m = \frac{k_2 + k_3}{k_1}$

(10) $V_{max} = k_3[E_T]$

(52) $K_i = \frac{k_6}{k_5} = \frac{[E][I]}{[EI]}$

(64) $[S_T] = [S] + [ES] + [ESI]$

(77) $\alpha K_i = \frac{k_8}{k_7} = \frac{[ES][I]}{[ESI]}$

(78) $[E_T] = [E] + [EI] + [ES] + [ESI]$

(79) $\frac{d[ES]}{dt} = 0 = [E][S]k_1 + [ESI]k_8 - [ES]k_2 - [ES]k_3 - [ES][I]k_7$

From (52) we obtain (80) $[EI] = \frac{[E][I]k_5}{k_6}$, and from (77) we get (81) $[ESI] = \frac{[ES][I]k_7}{k_8}$

Substituting these into (78) we obtain:

(82) $[E_T]=[E]\left(1+\frac{[I]k_5}{k_6}\right)+[ES]\left(1+\frac{[I]k_7}{k_8}\right)$

Solving (82) for [E]:

(83) $[E]=\frac{[E_T]-[ES]\left(1+\frac{[I]k_7}{k_8}\right)}{\left(1+\frac{[I]k_5}{k_6}\right)}$

On the other hand, substituting (81) into (64):

(84) $[S]=[S_T]-[ES]\left(1+\frac{[I]k_7}{k_8}\right)$

We can substitute (81), (83) and (84) into (79) and obtain: (85)

$$0=\left([E_T]\left(\frac{1}{1+\frac{[I]k_5}{k_6}}\right)-[ES]\left(\frac{1+\frac{[I]k_7}{k_8}}{1+\frac{[I]k_5}{k_6}}\right)\right)\left([S_T]-[ES]\left(1+\frac{[I]k_7}{k_8}\right)\right)k_1+[ES]\left(\frac{[I]k_7}{k_8}\right)k_8-[ES]k_2-[ES]k_3-[ES][I]k_7$$

Expanding the products and cancelling the last two terms in $k_7$, we obtain the expression (86)

$$0=k_1[E_T][S_T]\left(\frac{1}{1+\frac{[I]k_5}{k_6}}\right)-k_1[ES][E_T]\frac{\left(1+\frac{[I]k_7}{k_8}\right)}{\left(1+\frac{[I]k_5}{k_6}\right)}-k_1[ES][S_T]\frac{\left(1+\frac{[I]k_7}{k_8}\right)}{\left(1+\frac{[I]k_5}{k_6}\right)}+k_1[ES]^2\frac{\left(1+\frac{[I]k_7}{k_8}\right)^2}{\left(1+\frac{[I]k_5}{k_6}\right)}-[ES]k_2-[ES]k_3$$

After multiplyin both sides of the equality by $\frac{\left(1+\frac{[I]k_5}{k_6}\right)}{k_1\left(1+\frac{[I]k_7}{k_8}\right)^2}$ and gathering terms in [ES], it results in: (87)

$$0=[ES]^2-[ES]\frac{\left(1+\frac{[I]k_5}{k_6}\right)}{k_1\left(1+\frac{[I]k_7}{k_8}\right)^2}\left(k_1[E_T]\frac{\left(1+\frac{[I]k_7}{k_8}\right)}{\left(1+\frac{[I]k_5}{k_6}\right)}+k_1[S_T]\frac{\left(1+\frac{[I]k_7}{k_8}\right)}{\left(1+\frac{[I]k_5}{k_6}\right)}+k_2+k_3\right)+[E_T][S_T]\frac{\left(1+\frac{[I]k_5}{k_6}\right)}{k_1\left(1+\frac{[I]k_7}{k_8}\right)^2}\cdot\frac{k_1}{\left(1+\frac{[I]k_5}{k_6}\right)}$$

Using the continued fraction approach for the negative sign solution:

(88) $$[ES]=\frac{\dfrac{-\left(1+\dfrac{[I]k_5}{k_6}\right)}{k_1\left(1+\dfrac{[I]k_7}{k_8}\right)^2}[E_T][S_T]\cdot\dfrac{k_1}{\left(1+\dfrac{[I]k_5}{k_6}\right)}}{\dfrac{-\left(1+\dfrac{[I]k_5}{k_6}\right)}{k_1\left(1+\dfrac{[I]k_7}{k_8}\right)^2}\left(k_1[E_T]\dfrac{\left(1+\dfrac{[I]k_7}{k_8}\right)}{\left(1+\dfrac{[I]k_5}{k_6}\right)}+k_1[S_T]\dfrac{\left(1+\dfrac{[I]k_7}{k_8}\right)}{\left(1+\dfrac{[I]k_5}{k_6}\right)}+k_2+k_3\right)-[ES]}$$

Assuming [ES] << rest of the denominator, we can neglect it and also cancel the coefficients $\dfrac{-\left(1+\dfrac{[I]k_5}{k_6}\right)}{k_1\left(1+\dfrac{[I]k_7}{k_8}\right)^2}$. To simplify, we then multiply numerator and denominator by

$$\frac{\left(1+\dfrac{[I]k_5}{k_6}\right)}{k_1}$$

(89) $$[ES]\simeq\frac{[E_T][S_T]}{[E_T]\left(1+\dfrac{[I]k_7}{k_8}\right)+[S_T]\left(1+\dfrac{[I]k_7}{k_8}\right)+\left(\dfrac{k_2+k_3}{k_1}\right)\left(1+\dfrac{[I]k_5}{k_6}\right)}$$

Substituting (77) into (4), grouping terms, and using the definitions of the macroscopic kinetic constants (5), (10), (52), (65), we finally obtain the rate law:

(90) $$\frac{d[P]}{dt}=\frac{V_{max}[S_T]}{\left([E_T]+[S_T]\right)\left(1+\dfrac{[I]}{\alpha K_i}\right)+K_m\left(1+\dfrac{[I]}{K_i}\right)}$$

**(D) General case of activation/inhibition of a monomolecular irreversible reaction (linear and non-linear cases), under QSS assumption.**

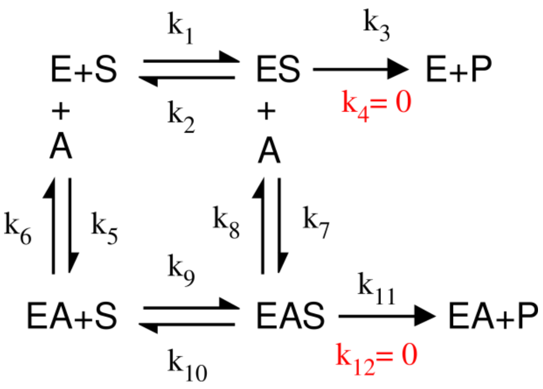


Similarly to the previous cases, a single substrate, irreversible reaction, in the presence of an effector (A) that could affect enzyme action either as an activator or as an inhibitor, being its effect partial or total, will be described by the following expressions:

Assuming [A] >> [$E_T$].

(5) $K_m = \dfrac{k_2 + k_3}{k_1}$

(10) $V_{max} = k_3[E_T]$

(91) $[E_T] = [E] + [EA] + [ES] + [ESA]$

(92) $[S_T] = [S] + [ES] + [ESA]$

(93) $\dfrac{d[ES]}{dt} = 0 = [E][S]k_1 + [ESA]k_8 - [ES]k_2 - [ES]k_3 - [ES][I]k_7$

(94) $\dfrac{d[P]}{dt} = k_3[ES] + k_{11}[ESA]$

(95) $K_a = \dfrac{k_6}{k_5} = \dfrac{[E][A]}{[EA]}$

(96) $\alpha K_a = \dfrac{k_8}{k_7} = \dfrac{[ES][A]}{[ESA]}$

(97) $k_{11} = \beta k_3$

Due to thermodynamic constraints, (98) $\alpha K_m = \dfrac{k_{10} + k_{11}}{k_9}$

From expression (95) we obtain (99) $[EA] = [E]\dfrac{[A]k_5}{k_6}$, and from (96) we can write

(100) $\left[EAS\right]=\left[ES\right]\frac{\left[A\right]k_7}{k_8}$

Substituting expressions (99) and (100) into the enzyme mass conservation expression (91) we get:

(101) $\left[E_T\right]=\left[E\right]\left(1+\frac{\left[A\right]k_5}{k_6}\right)+\left[ES\right]\left(1+\frac{\left[A\right]k_7}{k_8}\right)$ from which we can obtain [E]:

(102) $\left[E\right]=\frac{\left[E_T\right]-\left[ES\right]\left(1+\frac{\left[A\right]k_7}{k_8}\right)}{1+\frac{\left[A\right]k_5}{k_6}}$

Also, using (84) into (80) and rearranging for [S]:

(103) $\left[S\right]=\left[S_T\right]-\left[ES\right]\left(1+\frac{\left[A\right]k_7}{k_8}\right)$

Substituting expressions (100), (102) and (103) into the differential expression (93) we obtain: (104)

$$0=\left(\frac{\left[E_T\right]-\left[ES\right]\left(1+\frac{\left[A\right]k_7}{k_8}\right)}{1+\frac{\left[A\right]k_5}{k_6}}\right)\left(\left[S_T\right]-\left[ES\right]\left(1+\frac{\left[A\right]k_7}{k_8}\right)\right)k_1+\left[ES\right]\frac{\left[A\right]k_7}{k_8}k_8-\left[ES\right]k_2-\left[ES\right]k_3-\left[ES\right]\left[A\right]k_7$$

After cancelling the last two terms in $k_7$ and expanding, we reach: (105)

$$0=\frac{\left[E_T\right]\left[S_T\right]k_1}{1+\frac{\left[A\right]k_5}{k_6}}-\frac{\left[E_T\right]\left[ES\right]k_1\left(1+\frac{\left[A\right]k_7}{k_8}\right)}{\left(1+\frac{\left[A\right]k_5}{k_6}\right)}-\frac{\left[S_T\right]\left[ES\right]k_1\left(1+\frac{\left[A\right]k_7}{k_8}\right)}{\left(1+\frac{\left[A\right]k_5}{k_6}\right)}+\frac{\left[ES\right]^2k_1\left(1+\frac{\left[A\right]k_7}{k_8}\right)^2}{\left(1+\frac{\left[A\right]k_5}{k_6}\right)}-\left[ES\right]k_2-\left[ES\right]k_3$$

We can simplify by multiplying both sides of the equality by $\frac{1+\frac{\left[A\right]k_5}{k_6}}{k_1}$, so we obtain: (106)

$$0=\left[E_T\right]\left[S_T\right]-\left[E_T\right]\left[ES\right]\left(1+\frac{\left[A\right]k_7}{k_8}\right)-\left[S_T\right]\left[ES\right]\left(1+\frac{\left[A\right]k_7}{k_8}\right)+\left[ES\right]^2k_1\left(1+\frac{\left[A\right]k_7}{k_8}\right)^2-\left[ES\right]\frac{k_2+k_3}{k_1}\left(1+\frac{\left[A\right]k_5}{k_6}\right)$$

Gathering terms in [ES] and dividing both sides by $\left(1+\frac{\left[A\right]k_7}{k_8}\right)^2$

(107) $$0=[E_T][S_T]\frac{1}{\left(\frac{[A]k_7}{k_8}\right)^2}-[ES]\left(\frac{1}{\left(\frac{[A]k_7}{k_8}\right)^2}\right)\left(([E_T]+[S_T])\left(1+\frac{[A]k_7}{k_8}\right)+\frac{k_2+k_3}{k_1}\left(1+\frac{[A]k_5}{k_6}\right)\right)+[ES]^2$$

Using the continued fraction approach for the negative sign solution:

(108) $$[ES]=\frac{[E_T][S_T]\frac{1}{\left(\frac{[A]k_7}{k_8}\right)^2}}{\left(\frac{1}{\left(\frac{[A]k_7}{k_8}\right)^2}\right)\left(([E_T]+[S_T])\left(1+\frac{[A]k_7}{k_8}\right)+\frac{k_2+k_3}{k_1}\left(1+\frac{[A]k_5}{k_6}\right)\right)-[ES]}$$

Assuming [ES] << rest of the denominator, we can cancel it and also the coefficients

$$\frac{1}{\left(1+\frac{[A]k_7}{k_8}\right)^2}$$

(109) $$[ES]\simeq\frac{[E_T][S_T]}{([E_T]+[S_T])\left(1+\frac{[A]k_7}{k_8}\right)+\frac{k_2+k_3}{k_1}\left(1+\frac{[A]k_5}{k_6}\right)}$$

From (100) and (109):

(110): $$[EAS]\simeq\frac{[E_T][S_T]\frac{[A]k_7}{k_8}}{([E_T]+[S_T])\left(1+\frac{[A]k_7}{k_8}\right)+\frac{k_2+k_3}{k_1}\left(1+\frac{[A]k_5}{k_6}\right)}$$

Substituting (97), (109) and (110) into (94):

(111) $$\frac{d[P]}{dt}=\frac{[E_T][S_T]k_3+[E_T][S_T]\beta k_3\frac{[A]k_7}{k_8}}{([E_T]+[S_T])\left(1+\frac{[A]k_7}{k_8}\right)+\frac{k_2+k_3}{k_1}\left(1+\frac{[A]k_5}{k_6}\right)}$$

Gathering terms and substituting microscopic constants by their macroscopic counterparts using definitions (5), (10), (95) and (96), we obtain the rate law:

(112) $$\frac{d[P]}{dt}=\frac{V_{max}[S_T]\left(1+\frac{\beta[A]}{\alpha K_a}\right)}{([E_T]+[S_T])\left(1+\frac{[A]}{\alpha K_a}\right)+K_m\left(1+\frac{[A]}{K_a}\right)}$$
